# An *Ex Vivo* Muscle Physiology Method for Robust Measurement of Supraspinatus Muscle Function in Mouse Models

**DOI:** 10.64898/2026.01.12.698992

**Authors:** Baylah Mazonson, Hana Kalčo, Paola Divieti Pajevic, LaDora V. Thompson, Brianne K. Connizzo

## Abstract

The supraspinatus is the most frequently injured rotator cuff muscle, but its anatomical characteristics such as larger size, complex fiber architecture, and a single exposed tendon have limited the development of reproducible *ex vivo* contractility assays. In this study, we establish a robust method for *ex vivo* assessment of murine supraspinatus contractile function and characterize its physiological properties across age and injury conditions. We additionally adapt a barium chloride (BaCl_2_)–induced injury protocol for the supraspinatus, an approach not previously described, to evaluate how acute myofiber degeneration affects muscle performance. Male C57BL/6 mice (4 months) underwent 1.2% BaCl_2_ injection directly into the supraspinatus to induce controlled myofiber necrosis, allowing comparison of contractile behavior between injured and uninjured muscles. Using our injury *ex vivo* physiological testing protocol, we quantified optimal length (L_0_), twitch kinetics, force–frequency responses, peak tetanic force, and preliminary fatigue–recovery dynamics. Our protocol consistently generated fused tetanic contractions and reproducible force–frequency curves in the supraspinatus. We observed differences in supraspinatus contractility between young and old mice, consistent with well-established age-related changes in hindlimb muscle contractility. In addition, BaCl_2_ injury produced significant impairments in contractility 48 hours post-injection, demonstrating the sensitivity of this method to acute muscle damage. This study provides a novel and reliable method for evaluating the contractile function of the murine supraspinatus muscle ex vivo, overcoming previous anatomical challenges.

## Introduction

Rotator cuff pathologies present one of the most common causes of shoulder pain and disability, affecting millions of individuals worldwide each year.^1^ This includes shoulder dislocations and frozen shoulder syndrome, as well as degeneration and tears in the rotator cuff tendons. These injuries are especially prevalent among older adults, overhead athletes, and individuals exposed to repetitive mechanical loading or acute trauma.^1 2,3^ The most commonly injured rotator cuff tendon is the supraspinatus tendon, which passes laterally beneath the acromion and over the humeral head, integrating with the glenohumeral joint capsule.^2^ Together with the infraspinatus, teres minor, and subscapularis, the supraspinatus contributes to the dynamic stabilization of the glenohumeral joint by maintaining the humeral head centered within the shallow glenoid fossa throughout shoulder motion.^2^ Despite advances in surgical repair techniques and rehabilitation protocols, supraspinatus re-tear rates remain alarmingly high, often exceeding 40–60% in certain populations.^3^ Patients with chronic or recurrent tears often present with muscle pathologies, such as fatty infiltration, atrophy, and/or fibrosis, leading to impaired function.^4^ This highlights a persistent gap in our understanding of rotator cuff muscle biology, particularly contractile dysfunction, fatigue susceptibility, and age-related decline, as well as the crosstalk between tendon and muscle.^2^

Muscle contractility is a sensitive and reliable indicator of muscle health, reflecting the integrated function of sarcomeric structure, calcium handling, neuromuscular integrity, and metabolic capacity. Standard physiological assessments, such as twitch kinetics, tetanic force generation, fatigue–recovery dynamics, and specific force calculations, provide essential insights into both baseline muscle performance and the impact of aging, injury, or disease. These assessments can be performed ***in vivo, in situ***, or ***ex vivo***, each with distinct advantages and limitations.^5,6^ Although contractile testing of the supraspinatus muscle has been reported,^7,8^ a reproducible, well-characterized *ex vivo* method has not been widely established. *Ex vivo* muscle physiology, where the muscle is isolated from the animal and tested under strictly controlled environmental conditions, eliminates neural confounders and allows for precise manipulation of temperature, oxygenation, and stimulation frequency. However, this technique is best suited to small, thin muscles with two exposed tendons, such as the extensor digitorum longus (EDL) and soleus, because their geometry facilitates adequate diffusion of oxygen and nutrients to the muscle core.

In contrast, the murine supraspinatus presents several technical challenges that have limited its use in *ex vivo* studies. First, the supraspinatus muscle is larger and possesses only a single exposed distal tendon, inserting onto the humeral head; the proximal attachment is embedded within the scapula. This makes securing the muscle to a force transducer and fixed lever arm nontrivial, as standard dual-tendon mounting techniques cannot be applied. Second, the thicker architecture of the supraspinatus, with its pennation angle and relatively short fibers, raises concerns about adequate perfusion of the muscle core during prolonged testing, risking hypoxia or premature rigor.^9,10^ These anatomical challenges are valid in human, as well as small and large animal models. Animal models have been indispensable for studying rotator cuff pathology, enabling controlled manipulation of injury severity, timing, and loading environment. In particular, the rodent rotator cuff has been extensively used to characterize rotator cuff disorders and their pathologies. However, the lack of a standardized assay to directly quantify supraspinatus muscle contractile physiology *ex vivo* constrains our ability to evaluate muscle-specific functional deficits, recovery dynamics, and responses to injury in the rotator cuff complex.

A standardized protocol would enable researchers to quantify baseline murine supraspinatus function, compare contractile properties across lifespan, and assess how damaging stimuli (such as chemical injury) alter force production and fatigue dynamics. For example, chemical injury using barium chloride (BaCl_2_) is a well-established and reproducible model for inducing acute skeletal muscle damage in rodents. Upon injection, BaCl_2_ selectively disrupts the sarcolemma, triggering rapid and uniform necrosis of myofibers while preserving satellite cells, vasculature, and the extracellular matrix scaffold.^11^ This generates a controlled injury that mimics key aspects of human muscle trauma, including fiber degeneration, inflammatory infiltration, and subsequent regeneration. To our knowledge, BaCl_2_ injury has not previously been adapted for the murine supraspinatus muscle; therefore, we established and optimized a supraspinatus-specific BaCl_2_ injection protocol for this study. This model enables precise assessment of how degeneration and early regenerative processes alter supraspinatus force production, twitch kinetics, and fatigue– recovery dynamics.

The present study establishes a reproducible *ex vivo* muscle physiology protocol for the murine supraspinatus and characterizes its contractile properties. We provide a detailed method for supraspinatus dissection, mounting, oxygenation, and stimulation, optimized to preserve tissue viability and ensure accurate force measurements. Using this approach, we assess optimal length (L_0_), peak twitch force and associated kinetics, force–frequency relationships, peak tetanic force and associated kinetics, and fatigue responses. To demonstrate the utility of the method in the context of aging and muscle injury, we compare supraspinatus contractile function across age groups and with BaCl_2_ induced muscle injury. Together, these results establish a foundation for systematic investigation of supraspinatus physiology in aging and injury, providing a new tool to advance the field of muscle physiology.

## Materials

### Animals

Male C57BL/6 mice 4 months of age (n=5), 13 months of age (n=5) and 24 months of age (n=5) were purchased from Jackson Laboratories and housed in a temperature-controlled (25°C) facility on a 12-hour light/dark cycle at the Boston University vivarium, with ad libitum access to standard rodent chow and water. All procedures were approved by the Boston University Institutional Animal Care and Use Committee (IACUC).

### Solutions

Krebs-Ringer buffer was prepared fresh daily by adding 115 mM NaCl, 5.9 mM KCl, 1.2 mM MgCl_2_, 1.2 mM NaH_2_PO_4_, 1.2 mM Na_2_SO_4_, 2.5 mM CaCl_2_, 25 mM NaHCO_3_, and 10 mM dextrose to distilled water and stirring until dissolved.^12,13^ All tissue baths were filled with Krebs-Ringer buffer and heated to 25°C. Baths were oxygenated with 95% oxygen and 5% carbon dioxide.

## Methods and Results

### Dissection and Muscle Preparation

Animals were anesthetized with isoflurane and maintained on a temperature-controlled heating pad throughout the procedure to preserve physiological temperature. As reported in figure 1, the supraspinatus region was exposed by removing the overlying skin (Figure 1A.1) and superficial musculature (Figure 1A.2). The scapula was transected inferior to the scapular spine and through the infraspinatus muscle, taking care to avoid mechanical disruption of the supraspinatus (Figure 1A.3). The humerus was severed distal to the humeral head, and the clavicle was transected distal to the acromioclavicular joint. The supraspinatus, along with its associated scapular segment and humeral head, was immediately transferred to an oxygenated Krebs–Ringer tissue bath at a stable 25°C. Care was taken to avoid damaging the subclavian artery and to minimize the interval during which the supraspinatus was deprived of perfusion prior to immersion in the oxygenated bath. After both supraspinatus muscles were harvested, euthanasia was completed by cervical dislocation. In the tissue bath, the supraspinatus muscle was further refined by carefully removing residual tissues while preserving the integrity of the muscle belly and tendon. Remaining rotator cuff muscles, both infraspinatus and subscapularis, were mechanically disrupted (‘shredded’) using fine forceps to eliminate any potential contribution to force generation, ensuring that all contractile measurements reflected supraspinatus-specific function.

**Fig. 1.**
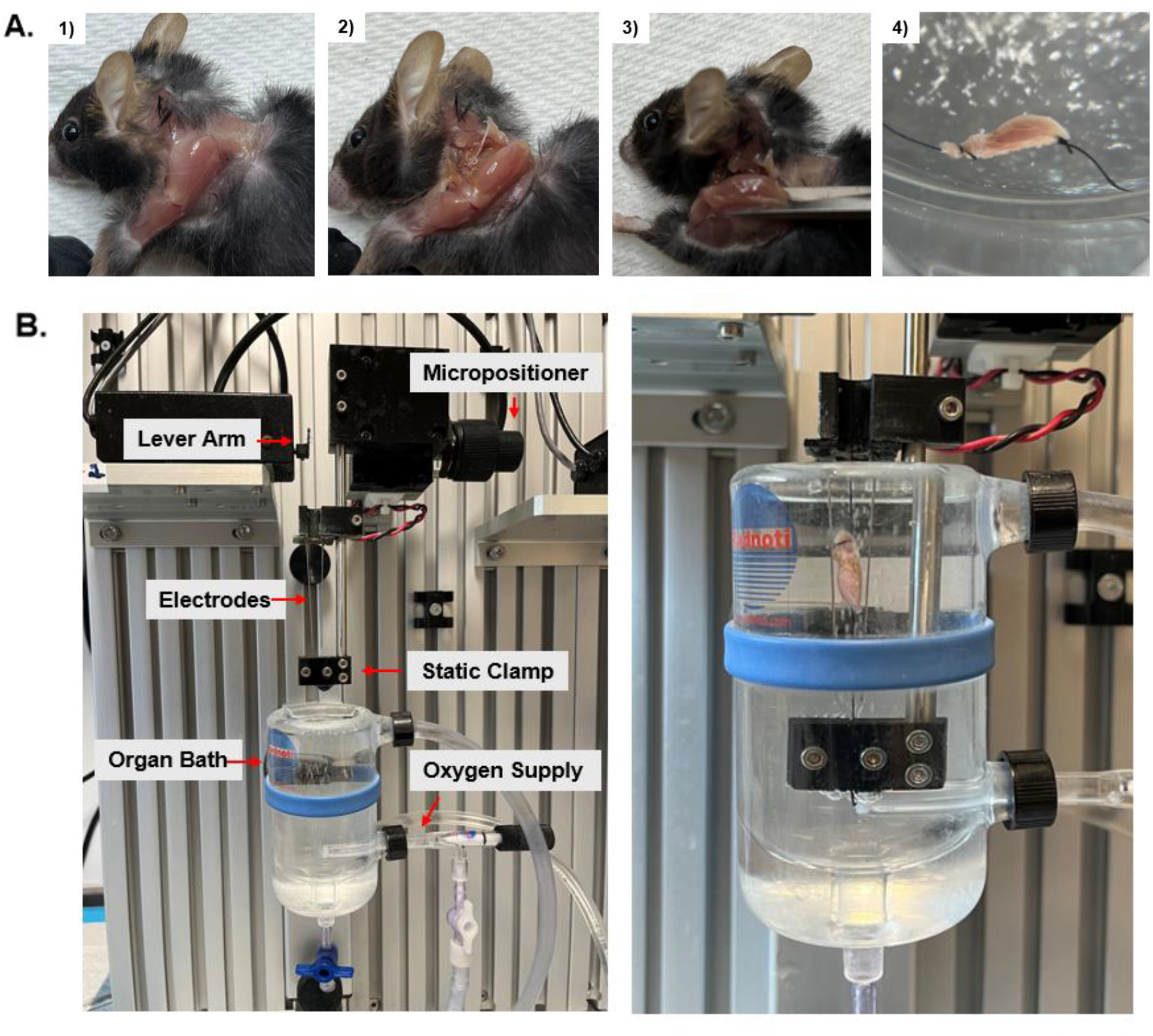
Supraspinatus isolation and *ex vivo* setup. A. Dissection of the supraspinatus. A.1) removing the superficial skin; A.2) removing superficial muscle; A.3) isolating the muscle by severing the scapula under the scapular spine and through the humerus; and A.4) tying around the humoral head and through the scapula once the supraspinatus has been transferred to an oxygenated organ bath. B. The *ex vivo* muscle physiology testing apparatus with and without the supraspinatus muscle secured. The muscle is secured with the proximal end to the static clamp and the distal end to the lever arm, suspended between the two electrodes in the oxygenated water bath. The length and preload tension of the muscle is adjusted with the micropositioner, which moves the static clamp vertically, thus increasing the tension on the muscle to hold it at optimal length.

Next, the bone-muscle-tendon–complex was secured at both ends using 4-0 braided silk suture. For proximal fixation, a small perforation was created in the scapular segment immediately inferior to the supraspinatus fossa using fine-point forceps, through which the suture was threaded and tied to provide stable anchorage without compromising bone integrity. Distal fixation was achieved by tying the suture around the humoral head, ensuring consistent alignment of the muscle–tendon unit (Figure 1A.4). The bone-muscle-tendon–complex was then mounted in the tissue bath, positioned longitudinally with the muscle belly centered equidistant between two platinum stimulating electrodes while avoiding direct electrode contact. The distal suture was connected to a calibrated force transducer, and the proximal scapular segment was secured to a static clamp to maintain constant muscle length during contractility testing (Figure 1B).

### Muscle Contractility

Muscle contractility was measured using a dual-bath physiology system (Aurora Scientific, Aurora, Ontario, Canada) consisting of two force transducers (0.5 N, Model 300B), two stimulators (Model 701B), one Dual Lever A/D Interface (Model 604B), one Dual System Signal Interface, customized Aurora software Dynamic Muscle Control (version 4.1.4.6), Dynamic Muscle Analysis (version 3.2), and a temperature control with water bath unit (Model 912; Polyscience, Niles, IL). Stimulation was delivered in biphase modality at an output of 1000 mA and 30 V, controlled by a dedicated Windows-based computer.

Muscle contractility measures were conducted using a well-established protocol for the EDL with modifications to preserve supraspinatus tissue viability. ^8,12^ As reported in Figure 2A, optimal length (*L*_0_) and maximal twitch force (*P*_t_) were determined by subjecting the supraspinatus to single-pulse square wave stimulations while holding the muscle at increasing lengths, which concurrently increased baseline (preload) force. *P*_t_ and *L*_0_ were measured at the start of the plateau of force production, defined as the first point where increasing the preload tension did not increase the maximum force produced by a single pulse stimulation. Increasing the preload beyond this point produced contractions of a consistent magnitude. At this point, *L*_0_ was measured with calipers from the supraspinatus tendon to the bottom of the scapula. All analyses were performed on GraphPad Prism (version 10.5), and significance set at P < 0.05. All other data are expressed as mean ± SEM. For reference, male mice weighing 27-33 grams, *L*_0_ was 10.33 ± 0.07 mm and *P*_t_ was 77.29 ± 5.99 mN. The muscle was maintained at *L*_0_ for all subsequent measures.

**Fig. 2.**
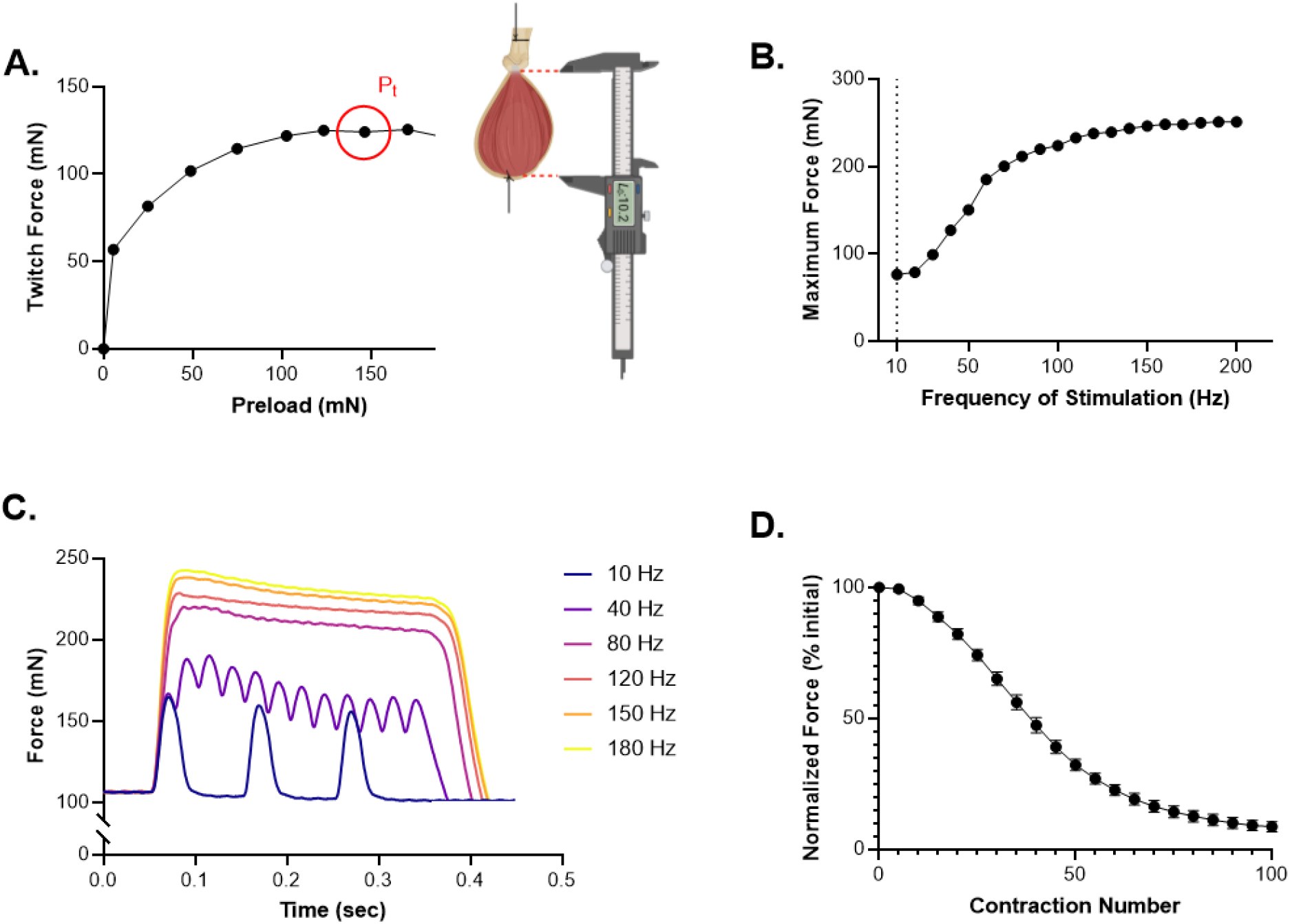
Representative experiments of supraspinatus to determine optimal length (*L*_o_), maximal twitch force (*P*_t_), force-frequency relationship, and peak tetanic contraction (P_o_). A. Force generating capacity with single-twitch stimulation over increasing preload forces (5-200 mN). *P*_t_ (red circle; 124 mN) occurred at 146 mN of preload. *L*_o_ (10.20 mm) was determined with calipers at *P*_t_. B. Force-frequency curve starting at 10Hz and increasing to 200 Hz, used to identify P_o_. P_o_ is achieved at 150 Hz and maintained through 200 Hz. C. Individual force recordings at submaximal and maximal frequencies, with a preload of 110 mN. D. Force production expressed as a percentage of initial force is significantly reduced when the supraspinatus is stimulated to fatigue at 80 Hz for 100 contractions, 2 seconds apart (*P<* 0.0001).

To ensure accurate force measurements, the next contractile parameter is peak tetanic contractile force (P_o_). As reported in Figure 2B and 2C, P_o_ was investigated using a force-frequency curve in which the muscle was stimulated with increasing frequency stimulation every minute. Increasing the stimulation frequency produced fused contractions around 80 Hz, although stimulating at greater frequencies continued to increase force production. The maximum force plateaued between 150 Hz and 180 Hz. Force production increased minimally when stimulating at frequencies above 180 Hz, hence the frequencies 10, 40, 80, 120, 150 and 180 Hz were selected to generate force-frequency relationships. Utilizing these six frequencies provides a reliable measure of submaximal and maximal contraction. P_o_ was generally observed at 180 Hz and was 256.12 ± 23.34 mN in male mice weighing 27-33 grams.

The next contractile parameter, the fatigability of the supraspinatus muscle, was determined using a modified established EDL fatigue protocol.^12^ As referenced in Figure 2D, the muscle was subjected to 100 stimulations at 80 Hz (the lowest frequency required to produce a fused contraction), with two seconds between stimulations. To evaluate fatigue over time, one-way ANOVA was conducted. In male mice weighing 27-33 grams, after 25 contractions, mean force declined 28% relative to the first contraction. At 50 contractions, mean force declined 58%, and at 100 contractions, mean force declined 87%.

At the end of muscle contractility testing, the supraspinatus was isolated by sliding forceps under the tendon, cutting the tendon, then separating the muscle from the scapula. This was performed to determine specific tetanic force (force generating capacity normalized to muscle mass) and requires the calculation of physiological cross-sectional area (PCSA). To calculate PCSA we used the equation: muscle mass (g) / [*L*_0_ (cm) * 1.06 (g/cm^3^)]. P_o_ was then divided by PCSA to calculate specific tetanic force. Male mice weighing 27-33 grams had supraspinatus muscles that weighed 28.21 ± 0.88 mg and a mean specific tetanic force of 73.71 ± 9.09 mN/cm^2^.

### Contractility and Barium Chloride Injection

To induce muscle damage, 4-month-old mice (n=5) were injected with Barium Chloride in one supraspinatus muscle and saline in the other. Mice were anesthetized with isoflurane and positioned prone on a heated surgical platform. The shoulder was gently externally rotated to expose the supraspinatus region and bring the muscle belly closer to the skin surface. The overlying fur was shaved, and the skin was disinfected with alternating scrubs of 70% ethanol and povidone– iodine. The supraspinatus muscle was localized percutaneously by palpating two external anatomical landmarks: (1) the spine of the scapula, which runs horizontally across the dorsal shoulder, and (2) the superficial arterial landmark over the humeral head. The midpoint between these landmarks corresponds to the thickest portion of the supraspinatus muscle belly.

A Hamilton syringe fitted with a 32G needle was used to inject 5 μL of 1.2% BaCl_2_ (intramuscular injection). The needle was inserted at a shallow angle (∼30° relative to the skin surface) and advanced approximately 1–2 mm into the muscle. The solution was delivered slowly over ∼5 seconds, and the needle was held in place for an additional 5 seconds to minimize reflux. Sham controls received an identical volume of sterile saline. No incisions were made at any point. Mice recovered on a heated pad and were monitored daily for normal activity and limb use.

Muscles were collected 48 hours post-injection for *ex vivo* contractility assessment. *P*_t_ and P_o_ contractions, rate of force development (RFD), and half relaxation time were assessed. RFD was defined as the greatest rate of force increase over time during muscle contraction. For *P*_t_ contractions, half relaxation time was defined as the time it took the muscle to relax halfway to baseline from the peak force production. For P_o_ contractions, since the peak force production can occur early or late into the contraction, half relaxation time was measured as the time from the end of stimulation that the muscle took to relax halfway to baseline. Paired t-tests were used to compare *P*_t_ and P_o_ contractions (maximum force, RFD, and half relaxation time) between BaCl_2_ and control injected muscles.

Barium chloride injections significantly reduced maximum force production in both *P*_t_ and P_o_ contractions (Figure 3). RFD was also significantly reduced in muscles injected with barium chloride. For *P*_t_ contractions (Figure 3A), there was a 61.64% decrease in mean maximum force and a 61.53% decrease in RFD in BaCl_2_ injected muscles compared to saline injected muscles. For P_o_ contractions (Figure 3B), there was a 49.46% decrease in mean maximum force and a 59.84% decrease in RFD in BaCl_2_ injected muscles compared to saline injected muscles. Half relaxation time was not significantly altered in *P*_t_ or P_o_ contractions.

**Fig. 3.**
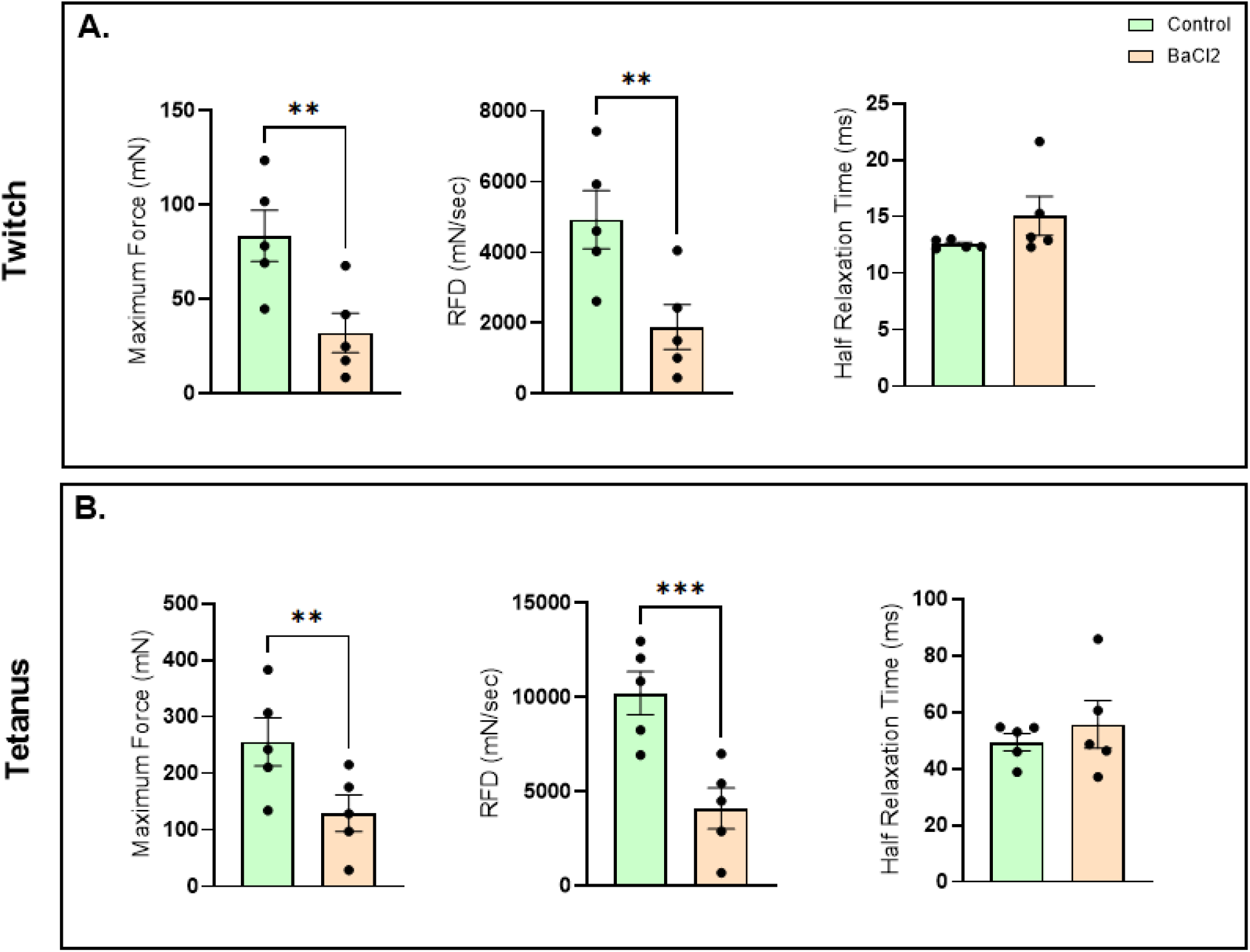
Barium chloride-related changes in supraspinatus contractility. 4-month-old mice (n=6) were injected with Barium Chloride (BaCl_2_) in one forelimb and saline in the other. Supraspinatus contractility was assessed 48 hours post-injection. BaCl_2_ reduced maximum force and rate of force development (RFD) compared to saline injected controls in 4-month-old mice in both P_t_ and P_o_. A. BaCl_2_ injection reduced maximum force and RFD during a single pulse twitch stimulation. B. BaCl_2_ injection reduced maximum force production and RFD during tetanic contractions. Half relaxation time was not significantly altered. \**P* < 0.05, \*\**P* < 0.01, \*\*\**P* < 0.001

### Contractility with Age

Mice aged 13 months (n=5) and 24 months (n=5) were compared to 4-month-old mice (n=5). The same contractile parameters were repeated from the barium chloride experiment. A one-way-ANOVA was used to compare single twitch and tetanic contractions between age groups. Where an interaction was detected, Tukey’s multiple comparisons tests were conducted.

Four-month-old mice displayed significantly greater force production and RFD during both *P*_t_ and P_o_ contractions than 13-month-old and 24-month-old mice (Figure 4). For *P*_t_ contractions (Figure 4A), there was a 55.20% decrease in mean maximum force and a 55.16% decrease in RFD in 24-month-old mice compared to 4-month-old mice. The time to half relaxation was not significantly different between age groups.

**Fig. 4.**
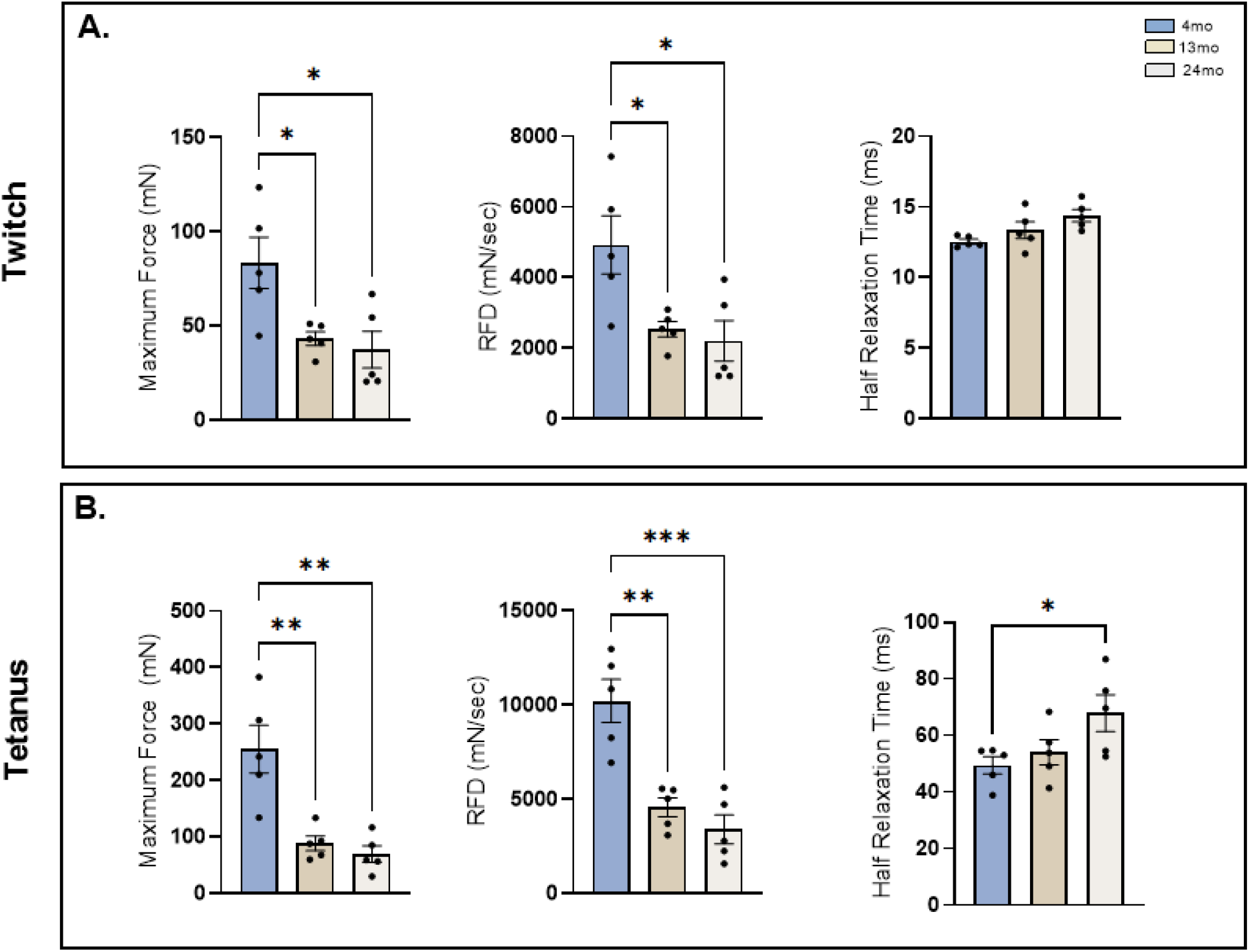
Age-related changes in supraspinatus contractility between 4 (n=6), 13 (n=6), and 24-month-old mice (n=5). Age reduces maximum force and rate of force development (RDF) in both P_t_ and P_o_. A. Single-pulse twitch maximum force and RDF are significantly lower in 13- and 24-month-old mice compared to 4-month-old mice. Time to half relaxation was not significantly different between age groups. B. Tetanic maximum force and RDF are significantly lower in 13- and 24-month-old mice compared to 4-month-old mice. Time to half relaxation was significantly greater in 24-than 4-month-old mice. \**P* < 0.05, \*\**P* < 0.01, \*\*\**P* < 0.001, \*\*\*\**P* < 0.0001

For P_o_ contractions (Figure 4B), there was a 72.66% decrease in mean maximum force and a 66.74% decrease in RFD in 24-month-old mice compared to 4-month-old mice. In addition, the time to half relaxation increased by 14.69% in 24-month-old mice from 4-month-old mice.

## Discussion

Rotator cuff injuries involve multiple soft tissues, but the supraspinatus muscle is particularly vulnerable to degeneration due to the high prevalence of tendon injury and/or chronic overload.^14^ The architectural complexity of the supraspinatus further underscores the need for dedicated physiological characterization. Rodent supraspinatus fibers are relatively short and arranged at a modest pennation angle, similar in organization to hindlimb muscles such as the semimembranosus, but adapted for stabilizing the glenohumeral joint, rather than forceful propulsion.^15,16^ This architecture enables fine-tuned force transmission across the rotator cuff but also makes the muscle particularly susceptible to alterations in tendon integrity, fiber atrophy, and fatty infiltration following injury.^15,16^ These features distinguish it mechanically from more commonly studied hindlimb muscles and highlight why direct physiological assessment of the supraspinatus, rather than extrapolation from limb muscles, is essential for understanding rotator cuff dysfunction. Measuring muscle contractility provides insight into multiple aspects of muscle health, including calcium handling, neuromuscular transmission, metabolic fatigue resistance, and the functional impact of structural damage. As a readout, contractile force is highly sensitive to subtle pathological changes and therefore serves as a valuable outcome measure for rotator cuff injury models.

Our findings demonstrate that the murine supraspinatus can reliably produce fused tetani at physiologically relevant stimulation frequencies and that its force-generating capacity can be quantified similarly to established hindlimb preparations. Our supraspinatus protocol was adapted from the EDL, a hindlimb muscle commonly used for *ex vivo* muscle physiology. The frequency at which maximal tetanic contraction (P_o_) is achieved is similar between muscles: 150 Hz for the EDL vs 180 Hz for the supraspinatus (Figure 2B and 2C). However, there are some notable differences between the two muscles. In the EDL, the preload necessary to generate P_o_ is on average between 15 mN and 25 mN. However, the preload necessary to generate P_o_ for the supraspinatus has a much larger range, generally falling between 50 mN and 150 mN (Figure 2A). The increase in and variability of required preload force for the supraspinatus is due to the pennate muscle architecture. In the EDL, there is a relatively uniform pennation angle and the fibers attach to one side of a single central tendon.^17^ In contrast, the structure of the supraspinatus is complex as fibers are short and pennate. These fiber and tendon arrangements in the supraspinatus influence the preload required to obtain optimal length (*L*_0_) and then generate P_o_.

Interestingly, the supraspinatus shows muscle-specific fatigue characteristics. How quickly a muscle loses force during repeated contractions is associated with the muscle’s metabolic capacity and calcium dynamics. In the current fatigue stimulation protocol, the point at which there is a 50% loss in force production occurs at the ∼25th contraction for the EDL and ∼40th contraction for the supraspinatus (Figure 2D). In contrast, the soleus muscle only fatigues by ∼35% after 100 contractions. The soleus shows a gradual decline in force or fatigue resistance, likely due to the muscle’s high oxidative capacity. The EDL shows a rapid decline in force or is highly fatigable in the presence of repeated contractions likely due to the muscle’s low oxidative capacity and reduced calcium release.^18^ The intermediate fatigue resistance characteristics found in the supraspinatus suggest a metabolic profile composed of both oxidative and glycolytic capacity, with greater glycolytic than oxidative capacity. Although supraspinatus fiber type is not well defined in mice, this metabolic profile is consistent with supraspinatus composition in rats.^19^

The supraspinatus muscle also exhibits the characteristic contractile alterations associated with aging and barium chloride injury. In aged mouse hindlimb muscles, isometric tetanic contractile force decreases, contraction time increases, and half relaxation time increases.^20,21^ Further, as an increase in time to peak contraction corresponds to a lower RFD, our results in the supraspinatus are consistent with well-established hindlimb contractility for muscles composed of type II fibers. Injection of BaCl_2_ into the supraspinatus muscle resulted in a significant reduction in maximum force generation and RFD compared to saline-injected controls in 4-month-old mice, as observed in both *P*_t_ and P_o_ contractions. The decrease in maximal force and RFD aligns with expected physiological consequences of acute muscle injury, which include loss of viable contractile tissue and alterations in muscle architecture. Notably, similar reductions in contractile force have been reported following BaCl_2_ injury in limb muscles such as the tibialis anterior and soleus, supporting the generalizability of these contractile deficits across different skeletal muscles.^22^

While our technique is an exciting step forward for the field, several limitations must be acknowledged. *Ex vivo* contractility inherently isolates the muscle from neural, vascular, and endocrine influences, which may limit its ability to fully recapitulate *vivo* physiology. Additionally, larger muscles such as the supraspinatus begin to exhibit early rigor or hypoxic decline if maintained in Krebs-Ringer buffer for extended durations; thus, only short or moderate fatigue protocols can be reliably executed. Future studies should explore improved perfusion strategies, including higher oxygenation flow rates or modified bath geometries, to extend tissue viability. Finally, while our bone-based anchoring method provides substantial stability, it may introduce slight variation in muscle alignment compared to tendon-only mounting. Despite these limitations, our findings establish a robust platform for investigating supraspinatus muscle physiology. This protocol will enable researchers to quantify functional deficits due to genetic modifications or injury, as well as adaptations to exercise and growth. Furthermore, pairing this approach with transcriptomic, proteomic, or imaging-based analyses could deepen our understanding of muscle degeneration and recovery in the context of shoulder injuries. In summary, we provide a technically accessible and physiologically sound method for *ex vivo* assessment of supraspinatus muscle contractility in mice. This tool fills a critical gap in rotator cuff research and can serve as a foundation for studying mechanisms of muscle dysfunction, testing therapeutic interventions, and modeling the complex interplay between muscle and tendon within the shoulder joint.

